# Choosing panels of genomics assays using submodular optimization

**DOI:** 10.1101/036137

**Authors:** Kai Wei, Maxwell W. Libbrecht, Jeffrey A. Bilmes, William Stafford Noble

## Abstract

Although the cost of high-throughput DNA sequencing continues to drop, extensively characterizing a given cell type using assays such as ChIP-seq and DNase-seq is still expensive. As a result, epigenomic characterization of a cell type is typically carried out using a small panel of assay types. Deciding a *priori* which assays to perform—e.g., a few complementary histone modification ChIP-seq experiments, perhaps an open chromatin assay, plus a few diverse transcription factor assays—is thus a critical step in many studies. Unfortunately, the field currently lacks a principled method for making these choices. We present submodular selection of assays (SSA), a method for choosing a diverse panel of genomic assays that leverages methods from the field of submodular optimization. We also describe a series of evaluation methods that allow us to measure the quality of a selected assay panel in the context of inference tasks such as data imputation, functional element prediction, and semi-automated genome annotation. Applying this evaluation framework to data from the ENCODE and Roadmap Epigenomics Consortia, we provide empirical evidence that SSA provides high quality panels of assays. The method is computationally efficient and is theoretically optimal under certain assumptions. SSA is extremely flexible, and can be employed to select assays for a new cell type or to select additional assays to be performed in a partially characterized cell type. More generally, this application serves as a model for how submodular optimization can be applied to other discrete problems in biology. SSA is available at http://melodi.ee.washington.edu/assay_panel_selection.html.

## Background

Genomics assays such as ChIP-seq, DNase-seq and RNA-seq can measure a wide variety of types of DNA activity, but the cost of these assays limits their application. In principle, to fully characterize a cell type, one would like to perform every possible type of assay, including ChIP-seq for a variety of histone modifications and dozens of transcription factors, several chromatin accessibility assays, and RNA-seq characterizing various types of RNA molecules. However, at current sequencing prices, performing a single genomics assay with reasonable sequencing depth costs on the order of $400 (https://www.scienceexchange.com/services/chip-seq. As a point of comparison, consider the ENCODE and Roadmap Epigenomics consortia, which develop, perform and analyze genomics assays as their primary activity (Bernstein et al., 2010; ENCODE Project Consortium, 2012). As of January 2015, the two consortia had performed a total of 216 types of assays on at least one cell type, and at least one assay on a total of 228 cell types (Methods). Applying all these assay types to all these cell types would require 49,248 assays; however, the two consortia have performed just 1,359 assays, 5% of the possible number (http://encodedcc.org; 2014). These consortia are worldwide efforts with large budgets; a typical lab might be able to perform at most several assays per cell type in order to analyze a particular tissue or perturbation that they are interested in. Moreover, there are virtually limitless perturbations and variations of a given cell type for which it would be interesting to examine the effect on DNA activity, including drug treatments, age, differentiation, etc.

Consequently, selecting a small panel of assays to perform on each cell type of interest—a problem we call *assay panel selection*—is a key step in any genomics project. To our knowledge, there has been little discussion in the literature of how to choose such a panel. In consortia such as ENCODE and Roadmap, the procedure for choosing which assay types to perform on each cell type is typically ad hoc. These decisions are made by the investigators involved, based on their intuition about the diversity of assay types, perhaps based on pairwise correlations between assays or similar simple metrics. Ernst and Kellis (Ernst and Kellis, 2015) proposed that imputation methods can evaluate the quality of a given of a given panel, but this approach cannot be used efficiently to select a panel (see Discussion).

In this work, we propose a principled method to solve the assay panel selection problem. Qualitatively, the method aims to identify, on the basis of existing data sets, assay types that yield complementary views of the genome. In practice, many pairs of assay types yield redundant information. For example, the transcription factors REST and RCOR1 are cofactors and therefore bind almost the same set of genomic positions (Andres et al., 1999). Similarly, the histone modification H3K36me3 primarily marks gene bodies, which are also transcribed and therefore measured by RNA-seq. Therefore, a great deal of what can be learned from the full set of assays can likely be learned by performing a small subset of the possible assays. This redundancy among assay types suggests that a carefully chosen panel of assays is likely to produce most of the information that would be obtained by performing all assays. Our solution to the assay panel selection problem is composed of two parts: an objective function that defines the quality of a panel, and an optimization algorithm that efficiently finds a panel that scores highly according to the objective function.

We propose to use an objective function called *facility location* (defined mathematically below), which measures what fraction of the information available in the full set of assay types is contained within the panel. This function has been previously applied in many fields, including document summarization (Lin and Bilmes, 2012), feature selection for machine learning (Liu et al., 2013), and exemplar-based clustering (Mirzasoleiman et al., 2013). This function also corresponds to the objective function of the widely-used k-medoids clustering algorithm (Gomes et al., 2010). Computing the facility location function requires a measure of similarity between assay types. For this purpose, we use the Pearson correlation between the two assays, averaged over the cell types in which the assays have been performed. We chose this objective function because it performs better than or comparably to the other methods we tried (Supplementary Notes 1−4).

To optimize the facility location function, we borrow methods from the field of submodular optimization. A simple approach to selecting a panel of assays would evaluate the facility location function for every possible subset of assays and then choose the highest-scoring subset. Unfortunately, 216 possible assay types yield 2^216^≈ 10^65^ possible panels of assays, so this approach is not feasible in practice. Fortunately, an efficient alternative selection method exists because the facility location function has the property of *submodularity.* The property of submodularity (defined mathematically below) is analogous to the property of convexity but is defined on discrete set functions rather than continuous functions. Submodular functions have a long history in economics (Vives, 2001; Carter, 2001), game theory (Topkis, 1998; Shapley, 1971), combinatorial optimization (Edmonds, 1970; Lovasz, 1983; Schrijver, 2004), electrical networks (Narayanan, 1997), operations research (Cornunejols et al., 1990), and more recently, machine learning (Narasimhan and Bilmes, 2005; Krause et al., 2008; Liu et al., 2013; Wei et al., 2014), but they are not yet widely used for problems in biology. Therefore, this application may serve as a model for how submodular optimization can be applied to biological problems more generally.

We apply existing submodular optimization algorithms to the facility location function to efficiently select a high-quality panel of assays, a method we call *submodular selection of assays* (SSA). There exists a large literature of methods for optimizing submodular functions. The optimization method we employ is very efficient and is theoretically guaranteed to find a solution whose quality comes within a constant factor of the quality of the optimal solution (Nemhauser et al., 1978a).

In addition to proposing solutions to the assay panel selection problem, an important contribution of this work is development of three general methods for evaluating the quality of a selected panel of assays. These three methods correspond to three distinct practical applications of the selected panel: (Equation 1) the accuracy with which the panel can be used to impute the results of assays not included in the panel; (Equation 2) the accuracy with which the panel can be used to detect functional elements such as transcription factor binding sites, promoters, and enhancers; and (Equation 3) the quality of a whole-genome annotation produced using the panel. These evaluation metrics share the property that an informative and diverse set of assay types yields better performance, according to each metric, than does a redundant set. Note that these evaluation metrics differ from the objective function because they use information that is not available at the time a panel is chosen; therefore, the evaluation metrics themselves cannot be used directly to choose a panel. These three metrics will be useful for any future study of the quality of a panel of assays, independent of the particular procedure used to choose such a panel.

We consider two variants of the assay panel selection problem. We are primarily interested in the “future” variant, which arises when a researcher is interested in applying a panel of genomics assays to a new tissue type or cellular condition. In this case, the researcher must use previously performed assays in other cell types to choose a representative panel of assay types. However, we also consider the “past” variant, which arises when a researcher is interested in applying a computationally expensive analysis, such as a genome annotation method, that cannot efficiently be run on all available data sets. In this setting, the researcher must choose a representative panel from the available data to use as input to the analysis. In this case, the researcher may use the data from assays performed on the cell type in question to inform their choice. We propose a variant of our method, called SSA-past, which leverages this information to allow a researcher to choose such a representative panel in this setting.

## Results

### Submodular Selection of Assays Identifies Diverse Panels of Genomics Assays

Submodular selection of assays (SSA) takes as input a collection of genomics data sets (“assays”) and identifies a high-quality subset of those assays (Methods, Figure 1). Each input data set is represented as a real-valued signal vector over the genome. SSA begins by computing a pairwise similarity matrix that contains, for each pair of assay types, the mean Pearson correlation over all pairs of assays of those two types. The method employs a submodular function, called *facility location* (Methods), to estimate the quality of any possible panel of assay types. The facility location function takes a high value for a particular panel when all assay types have at least one similar representative in the panel. We chose this strategy because it performed the best of those we tried according to our evaluation (Supplementary Notes 1-4). SSA then applies the *greedy submodular optimization* algorithm (Methods) to efficiently choose a panel of assays that maximizes this facility location function. The output of this method is an ordered list of assay types, where the top k assay types in this list represent a high-quality panel of size k. A detailed description and theoretical justification for the method is provided in Methods.

**Fig. 1.**
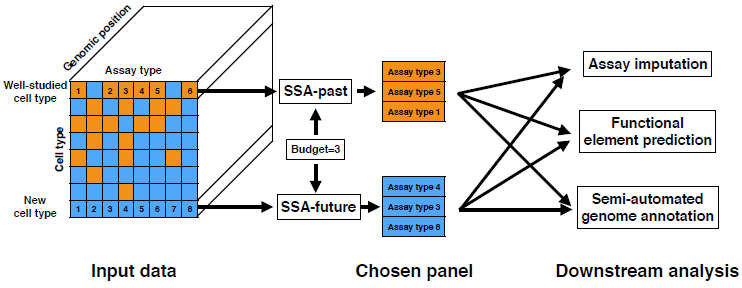
Schematic of the selection process performed by submodular selection of assays (SSA). The method takes as input all available existing genomics assays, where each assay is represented as a real-valued track over the genome. In the SSA-past mode, SSA selects a panel of already-performed assays to use as input to an expensive computational analysis. In the SSA-future mode, SSA chooses a panel of assay types to be performed in a new cell type. In both cases, the resulting data sets are provided as input to downstream analyses, which may include imputing assays that weren’t performed, predicting the locations of functional elements, or semi-automated genome annotation.

By analyzing data from the ENCODE and Roadmap Epigenomics Consortia, we found that SSA results in assay panels with diverse genomic functions. Because researchers generally perform panels of either histone modifications or transcription factor ChIP-seq assay types but rarely perform mixed panels, we ran the method separately on transcription factor and histone modification types (combined panels are shown in Supplementary Table 1). When choosing from transcription factors, SSA chooses factors that engage in diverse regulatory pathways (Figure 2). The vast majority of transcription factors in our data set bind to promoters and enhancers and regulate the transcription of RNA Pol II-transcribed genes. The top five transcription factors chosen by SSA include three of these factors, each of which regulate very different regulatory pathways: SMARCB1, an ATP-dependent chromatin remodeler; PML, a tumor suppression factor; and STAT5A, a factor involved in developmental signal transduction (UniProt Consortium, 2014). The top five also includes two factors—CTCF and BRF2—that are not solely involved in RNA Pol II-mediated transcription. CTCF, part of the cohesin complex, regulates chromatin conformation and enhancer-promoter insulation, and only about half of its binding sites occur in promoters or enhancers. BRF2 is part of the RNA Polymerase III complex, which transcribes rRNA, tRNA and other small RNAs. These two assay types each have low objective scores when in a panel by themselves (“singleton scores”), but are chosen by SSA because they measure different types of activity than the rest of the panel. Therefore, they are important to include in a diverse panel.

**Fig. 2.**
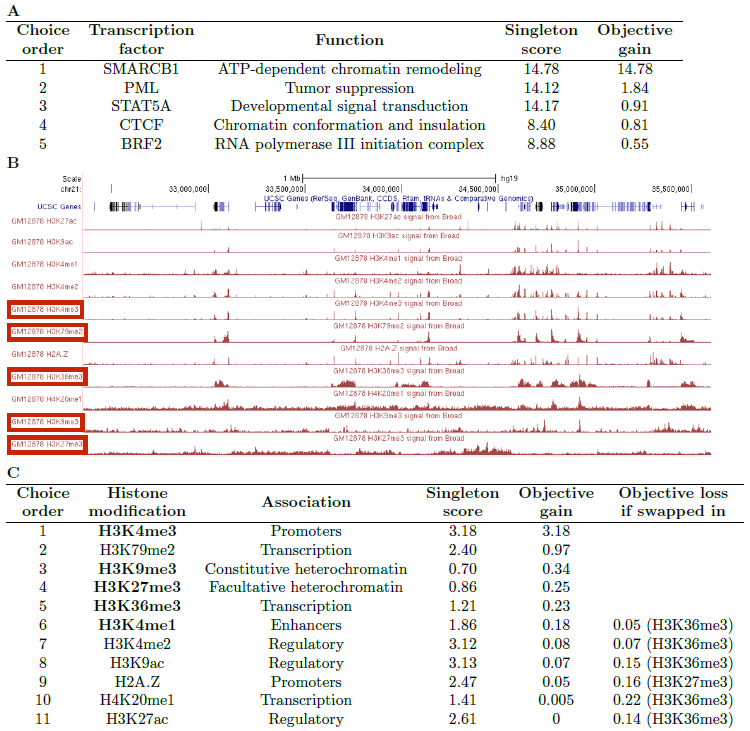
Results of SSA-past on ENCODE+Roadmap data sets. (A) Panels of transcription factors assays chosen by SSA-future. Each list is in the order assigned by SSA; for any size k, the top k assay types in the list are the chosen panel of this size. The “singleton score” is the objective value of a panel containing only the indicated assay type, and the “objective gain” indicates the improvement in objective that results from SSA adding the indicated assay type to the growing panel. Because there are 80 transcription factors, we display just the top five chosen by SSA. Associations are summarized from UniProt (UniProt Consortium, 2014). (B) Redundancy in histone modification signal in the genome. The top five assay types chosen by SSA are boxed in red. (C) Similar to (A), but for the histone modification assays. See the main text for a description of the “Objective loss if swapped in” column. There are only eleven histone modifications, so we display the full list. Bold font indicates those histone modification assays chosen by the Roadmap Epigenomics consortium.

When choosing a panel of histone modifications, SSA selects marks that cover diverse types of genomic regions and exhibit qualitatively different patterns ((Figure 2b),(Figure 2c). The top six histone modifications include a promoter mark (H3K4me3), an enhancer mark (H3K4me1), a gene mark (H3K79me2) and marks associated with both known types of repressive domains, facultative (H3K27me3) and constitutive (H3K9me3) heterochromatin. The two repressive marks, H3K27me3 and H3K9me3, have the lowest singleton scores of all the histone modifications, but they give a high objective gain because they measure distinct activity from the rest of the panel. In contrast, even though H3K27ac is sometimes considered the best individual mark of enhancers and has a high singleton score, it is chosen last by SSA because it is redundant with other assay types in the panel, such as H3K4me1 and H3K9ac. The top six includes two different marks of transcription, H3K79me2 and H3K36me3, but these two modifications mark different parts of genes and are regulated differently relative to the gene’s level of transcription (Li et al., 2007). As expected, SSA ranks additional measures of regulation (H3K4me2, H2A.Z, H3K9ac and H3K27ac) low on the list because thes marks are redundant with the regulatory marks H3K4me1 and H3K4me3.

SSA almost exactly recapitulates the panel of histone modifications chosen by the Roadmap Epigenomics consortium (boldface entries in Table 2C). This consortium chose a set of five “core” histone modifications to assay across 111 human primary tissues. This choice was made by the members of this consortium based on their collective, expert knowledge. These five core histone modifications ranked among the top six modifications chosen by SSA. In fact, the SSA-chosen and Roadmap-chosen panels of size 5 have very similar scores according to the facility location function, ranking 1 and 16 respectively out of all 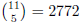 possible panels of five histone modifications. Therefore, SSA closely reproduces careful, manual selection by experts in an entirely automated and data-driven way.

To better understand the choices made by SSA in selection of histone modifications, we performed a “swap-out” experiment (final column of Table 2C). We started with the panel of size five selected via SSA, and we asked, for each of the remaining six histone modification assays, how much the objective function would decrease if we were forced to swap one of the SSA-selected assays for the excluded assay. In five out of the six cases, the objective is maximized by swapping the excluded assay with the last-selected histone modification, H3K36me3. However, the magnitude in the change in objective varies quite a bit: swapping in H3K4me1 makes very little difference (0.05), whereas swapping in H3K36me3 yields a relatively large change in objective (0.22). This type of exploratory analysis can be quite valuable in the context of a real experimental design setting, where qualitative features of the assays (e.g., familiarity to the researchers involved) are important but difficult to quantify.

In addition to its efficiency, SSA is flexible. First, in some circumstances, a researcher may have already performed a few assays in a given cell type or be confident that they want to perform them, and is therefore interested in choosing which assay types best complement these existing assays. SSA can be used in this scenario simply by restricting the returned set to include these assay types. For example, restricting the set of histone modifications to contain H3K27ac de-prioritizes other enhancer marks, such as H3K9ac and H3K4me1 (Supplementary Table 2). Second, a researcher might have a prior preference for certain classes of assays because they are easier to perform, less expensive, or measure features the researcher is especially interested in. SSA can include this prior preference by adding a weight to the objective for each assay type, reflecting the preference for that assay type. For example, selecting from all assay types (transcription factor, histone modification and DNA accessibility) jointly while placing a preference for (or against) histone modifications increases (or decreases) the fraction of histone modifications chosen to for the panel (Supplementary Table 3). Third, SSA can be used to select cell types instead of assay types by transposing the cell type-assay type matrix and running the same analysis. Doing so produces a panel of cell types that includes a diverse mixture of mesodermal, endodermal and undifferentiated cells, and both normal and cancer-derived cells (Supplementary Table 4).

### Three Metrics Evaluate the Quality of a Set of Genomics Assays

In order to quantitatively evaluate SSA, we developed an evaluation framework for assay panel selection. We focused on three of the most common downstream applications of genomics data sets: (Equation 1) imputing assays that have not been performed, (Equation 2) locating functional elements such as promoters and enhancers, and (Equation 3) annotating the genome using a semi-automated method. We describe each metric briefly here, with full details provided in Methods.

The first evaluation metric, *assay imputation*, measures how well a chosen panel of assays can be used to predict assays that have not been performed (Figure 3A). We train a regression model to predict each assay outside of the panel on the basis of the assays within the panel, using random subsets of the genome for training and testing, respectively. High performance on the assay imputation metric indicates that the panel contains all of the information in the assays outside of the panel. Moreover, recent work on imputation has showed that it is often effective to train a regression model on data from reference cell types and apply it to a target cell type (Discussion).

**Fig. 3.**
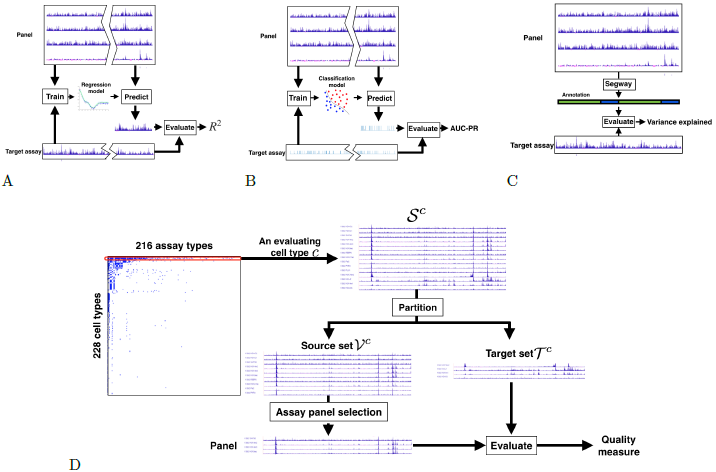
Evaluation strategies. Schematics of the three evaluation metrics: (a) assay imputation, (b) functional element prediction, and (c) annotation-based evaluation, as well as (d) the overall cross-validation evaluation strategy.

**Fig. 4.**
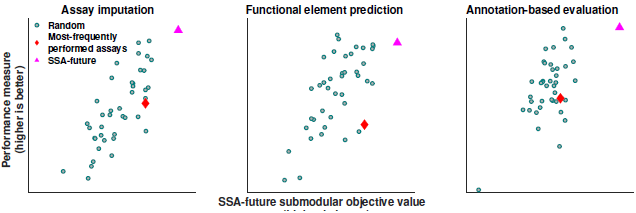
Relationship between the facility location objective function and evaluation metrics. Each dot corresponds to one of 40 randomly-chosen panels. Pink triangles indicate results from maximizing the SSA-future facility location function; red diamonds indicate the panel of most-frequently performed assay types (Supplementary Table 5). These results were computed in GM12878, using panels of four assay types.

The second evaluation metric, *functional element prediction*, is similar to the assay imputation metric but focuses specifically on how well a chosen panel of assays can be used to locate functional elements such as promoters and enhancers (Figure 3B). Because there are few validated examples of each type of element, we use experimentally determined binding of transcription factors, as measured by transcription factor ChIP-seq, as a proxy for functional elements. We train a classifier model to predict the locations of these elements on the basis of the assays within the panel. High performance on the functional element prediction metric indicates that a panel can be used to accurately locate functional elements. Although both the assay imputation and functional element prediction evaluation metrics aim to predict genomics data sets, functional element prediction focuses on the small fraction of the genome corresponding to transcription factor binding sites.

The third evaluation metric, *annotation-based evaluation,* measures how effectively a given panel can be used to annotate the genome through a semi-automated genome annotation (SAGA) method (Figure 3C). SAGA methods, which include HMMSeg (Day et al., 2007), ChromHMM (Ernst and Kellis, 2010), Segway (Hoffman et al., 2012) and others (Thurman et al., 2007; Lian et al., 2008; Filion et al., 2010; Kharchenko et al., 2010; Jaschek and Tanay, 2009; Larson et al., 2013; Biesinger et al., 2013), annotate the genome on the basis of a panel of genomics assays. They simultaneously partition the genome and annotate each segment with an integer label such that positions with the same label exhibit similar patterns of activity. These methods are semi-automated because a person must interpret the biological meaning of each integer label. SAGA methods have been shown to recapitulate known functional elements including genes, promoters and enhancers. Given a particular panel of assays, we perform annotation-based evaluation by using this panel as input to Segway and measuring how well the resulting genome annotation corresponds to patterns observed in the assays outside of the panel. High performance on this metric indicates that the chosen panel can be used to produce a comprehensive annotation of the genome.

Applying these metrics to evaluate a method for choosing panels is complicated by two factors. First, no cell type has had all assay types performed in it, so we perform evaluation separately on each cell type in order to evaluate against all available assay types. Second, these evaluation metrics must be used to compare to assays outside of the panel, so we use a cross-validation strategy in which we hold out a *target set* for evaluation and choose panels from the remaining *source set*, repeating this process for many choices of target set. This evaluation strategy enables principled evaluation that compares all methods against the same held-out standard while using only the available data sets (Methods, (Figure 3D).

### Panels Chosen by SSA Perform Well by Three Evaluation Metrics

We applied this panel evaluation framework to evaluate SSA. First, to determine the most effective objective function, we compared the facility location function and four other potential objective functions based on the pairwise similarity matrix. We found that the facility location function had a higher Spearman correlation with the three evaluation metrics than the other objective functions we tried, and this trend was consistent across multiple cell types (Supplementary Note 1). In addition, we found that the facility location function produced the highest correlation with the three evaluation metrics when we defined the similarity between a pair of assay types as the mean Pearson correlation between this pair, as opposed to the median, maximum or other aggregation function (Supplementary Note 2). We also compared between two strategies for deriving the Pearson correlation. The first computes the correlation based on random samples of genomic positions, and the second on the DNase peak positions only. We found that almost comparable performance is achieved on the three evaluation metrics for the two strategies (Supplementary Note 3). We lastly compared the effectiveness of using Pearson versus Spearman correlation for computing the similarity measure. We found that consistently better performance is achieved when the similarity is defined using the Pearson correlation (Supplementary Note 4). These observations led us to choose our variant of facility location as the SSA’s objective function.

Next, we used our three evaluation metrics to compare SSA to alternative panel selection approaches. As a baseline, we considered randomly selected panels of a given size. We also considered the panel of most frequently performed assays (Supplementary Table 5) as a good proxy for a likely data-driven choice. We found that the panels reported by SSA perform among the top few percent out of the space of all possible panels, and this high performance is consistent across panel sizes, evaluation cell types and performance metrics (Figure 5A). We found that SSA also greatly outperforms the panel of most-frequently performed assay types. Indeed, this commonly-performed panel actually performs worse than the average panel in many cases, which may be a consequence of the fact that the most commonly-performed assay types measure broad marks of regulation, such as histone modifications and DNA accessibility, which do not have the specificity to identify pathway-specific elements. These results demonstrate quantitatively that panels chosen by SSA are effective when applied to their most common downstream tasks.

**Fig. 5.**
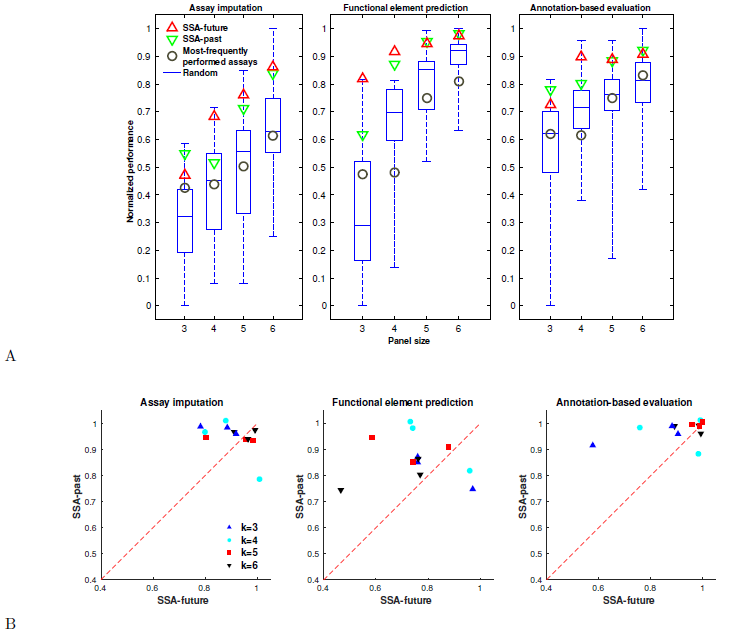
Performance of panel selection strategies. (a) Boxplots show the distribution of evaluation metrics over 40 random panels on data from cell type GM12878. The panels of most-frequently performed assays are composed of the top k most frequent assay types available in our data set, where k is the size of the panel. Each evaluation metric is normalized to lie within [0,1] by subtracting the lowest value and dividing by the highest. (b) Scatter plot between the performance of SSA-past and SSA-future across cell types K562, GM12878, and H1-hESC. Each dot in the plot corresponds to the two variants of SSA for a panel size evaluated using a metric in a cell type. The performance is measured as the fraction of panels that perform worse than the SSA-chosen panel, estimated by comparing to 40 randomly-selected panels.

### SSA can Select a Subset of Performed Assays as Input to an Expensive Analysis

So far we have considered panel selection in the “future” setting, where a researcher is planning to experimentally perform a panel of assay types. Panel selection is sometimes also important in the “past” setting, where a researcher wishes to apply a computationally expensive analysis that cannot be efficiently applied to all assays together and therefore must be applied to a smaller panel. For example, training a statistical model to perform semi-automated genome annotation jointly on dozens of assays across many cell types is computationally expensive. A strategy in which each cell type is represented by a smaller panel of assays might yield very similar annotations using a fraction of the computational resources. In this setting, the assays themselves are available to the selection algorithm, so we compute the similarity matrix based on these values themselves (SSA-past) rather than estimating the similarities by aggregating across cell types (SSA-future). Importantly, in the selection of past assays case, a different panel can be selected for each cell type, based on the available data. Alternatively, if the researcher would like to use the same panel of past assays for analyses several cell types, SSA-future can be used to select a panel that maximizes the average quality over all target cell types (Supplementary Table 6). To test SSA in the past setting, we used the same evaluation strategy as in the future setting, but using the source assays themselves to compute the similarity matrix. SSA performs consistently well according to these metrics, and it performs slightly better on some cell types in the past than the future setting due to the availability of this additional information (Figure 5B).

## Discussion

The growing availability of a large number of types of genomics assays means that choosing a panel of genomics assays is a key step in any genomics project. Previously, these panels were chosen in an ad hoc fashion. We have developed submodular selection of assays (SSA), a method for choosing high-quality panels using submodular optimization. This method is computationally efficient, results in high-quality panels according to several quality measures, and is mathematically optimal under some assumptions. By applying SSA, researchers can now easily choose a high-quality panel of assay types to perform on any cell type of interest. These higher-quality panels will allow researchers to achieve the same utility from performing fewer assays, saving thousands of dollars in labor and reagent costs per cell type. This panel selection framework can also be used partway through the investigation of a cell type, when several assays are already available. By modifying the facility location function to include the availability of these assays, SSA can be used to determine the most-informative next experiments to perform. In doing so, SSA will take into account the information in these existing assays and choose additional assay types that measure distinct genomic features.

A key feature of the submodular optimization approach is the flexibility afforded by the broad class of submodular objective functions, and the ability to encode appropriate prior knowledge into the selection of the objective. In this manuscript, we focused on optimizing the facility location function. However, the same submodular optimization framework can be used to optimize other objective functions that may prove to be more relevant for certain applications. Several other functions may be useful in practice. First, if some assays are more expensive or time consuming than others, then the objective function can be modified to incorporate this cost. Second, if some assays are inherently preferable to others, for example because they have better-established processing pipelines, then the objective can incorporate this preference and trade off choosing both diverse and established assay types. Third, entirely different types of panel attributes may be valuable for a particular application, which can be formalized as a different objective function, such as the alternatives we discuss in Supplementary Note 1. As long as the resulting objective function remains submodular, it will be efficiently optimizable using either the greedy algorithm for monotone non-decreasing functions or other efficient methods (Buchbinder et al., 2012, Buchbinder et al., 2014) for non-monotone functions. Moreover, such modifications are intuitive to design and easy to implement.

The facility location function can also be used to guide manual assay panel selection. A researcher may seek to optimize hard-to-quantify characteristics of a panel, such as familiarity with the protocols involved or the panel’s concordance with panels performed on other cell types. In this case, the researcher may choose to perform a panel that has slightly poorer quality as measured by the facility location function in order to optimize these other criteria. Still, in such a setting, manual investigation of the objective values associated with different panels can provide useful insights.

The framework presented here can also be extended to many related problems. We have discussed two such variants that apply in the scenarios, “I would like to select a panel of assay types to perform, taking cost into account”, and “I have carried out many assays in this cell type and would like to choose a subset of this to use as input to an expensive computational analysis”. This framework can also trivially be extended to the problem, “I have already performed several assay types in a particular cell type of interest and would like to select several more to perform,” by simply restricting the output panel to contain the previously-performed assays (Supplementary Table 2). The same framework also applies to the problem, “I have a set of assay types commonly performed in several cell types and would like to choose several assay types from the set that are the most informative and representative to study the cell types” by restricting the aggregation of the assay type similarity to the cell types at hand (Supplementary Table 6). In addition, by deriving a similarity measure between cell types, the methods presented here could be used to solve the problem, “Based on orthogonal data such as gene expression profiles, which new cell types should I perform assays in?” Finally, related methods may be able to apply to the problem “I have carried out assays across a variety of cell types and assay types, and I would like to select a set of additional assays to apply in any combination of cell and assay types.”

The evaluation strategy we introduce here can be used to evaluate any proposed strategy for panel selection, whether or not this method is based on submodular optimization. This framework is composed of two parts. First, a cross-validation strategy allows for principled comparison of methods under the restriction that not all assays are available in all cell types, and that a panel must not be evaluated on an assay type that it contains. Second, three distinct metrics capture the three primary downstream applications of genomics data sets.

The problem of assay panel selection was posed previously by Ernst and Kellis (Ernst and Kellis, 2015). These authors proposed the assay imputation evaluation metric for evaluating a panel of genomics assays. However, this type of evaluation metric cannot be used to choose an assay panel for two reasons. First, evaluating the accuracy of an imputation model requires that the target data set already be performed. Second, even if such data sets were available, evaluating all possible panels of size K from N assay types requires applying evaluation to 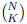 possible panels, which is hopelessly computationally expensive. For example, the method of Ernst and Kellis was reported to take roughly one hour to train per panel, so choosing a panel of five assay types from the 188 previously-performed assay types would take on the order of 10^5^ processor-years. In contrast, the facility location objective function we define does not require that the target data sets be available in order to evaluate the function, and the submodular optimization approach requires only a polynomial number of evaluations. Moreover, we define two additional evaluation metrics (functional element prediction and annotation-based evaluation) that complement evaluation by evaluation by assay imputation.

In addition the assay panel selection problem, the same submodular optimization framework may also be useful in selecting a set of informative and representative cell types to study. This cell type panel selection setting can be viewed as a dual variant of the assay panel selection problem (Supplementary Table 4). The flexibility of SSA (e.g., forcing selection of certain assay types, or weighing assay types based on cost or preference) easily carries over to the cell type selection setting. Moreover, the methodologies proposed here for evaluating the quality of a panel of assay types (e.g., assay imputation, and annotation-based evaluation) can be easily extended to assess the goodness a cell type panel selection approach.

One limitation of any data-driven analysis is that it is limited by any imperfections in the data sets used. For example, if all available assays of a given type happen to be of particularly good or poor quality, then the correlations associated with this assay type will appear to the algorithm to be particularly strong or weak, respectively. Similarly, any mislabeled assays, batch effects, or other artifacts may also influence whether certain assay types will be chosen in a panel. Future assays of that type may not be expected to exhibit the same artifactual patterns, so the resulting panels could be suboptimal. Therefore, it is always important to scrutinize the results of data-driven approaches like this one to understand whether patterns in the available data are predictive of future experiments. Modifications of this approach that, for example, find and remove faulty assays before input into the algorithm might result in different panels. However, our evaluation metrics are also entirely data-driven, so we cannot use them to explore these issues.

As noted above, submodular optimization is widely used for discrete problems in other fields but is not yet widely used in biology. We hope that the current work can serve as a model for how submodular optimization can be applied to other problems in biology. As with convex optimization, the same toolbox of submodular optimization methods can be applied to a wide variety of problems, and any innovations to this toolbox improve all solutions. Therefore, we expect that in the future, submodular optimization will be used for other discrete problems in biology, such as for selecting panels of DNA mutations to test in a functional screen or removing redundancy in protein sequence data sets.

## Methods

### Genomics Data

We acquired all public genomics data from the ENCODE (http://hgdownload.cse.ucsc.edu/goldenPath/hg19/encodeDCC/) and Roadmap Epigenomics (https://sites.google.com/site/anshulkundaje/projects/epigenomeroadmap) projects as of January 2015. These data sets were processed by the two consortia into real-valued data tracks, as described previously (Hoffman et al., 2013; Kundaje et al., 2015). We omitted all assays with more than 1% unspecified positions, which may indicate errors during processing or mapping. We manually curated these assays to unify assay type and cell type terminology and, when multiple assays were available, we arbitrarily chose a representative assay for each (cell type, assay type) pair. This procedure resulted in a total of 1,359 assays comprised of a total of 216 assay types and 228 cell types. The assay types include ChIP-seq with a variety of targets (both histone modification and transcription factor), DNase-seq, FAIRE-seq, Repli-seq and RNA-seq. The full list of assays is given as supplementary data. We applied the inverse hyperbolic sine transform asinh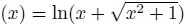 to all signal data. This function has the compressing effect of a function like log x for large values of x but it is defined at zero and has much less of a compressing effect for small values. The asinh transform has been shown to be important for reducing the effect of large values in analysis of genomics data sets (Johnson, 1949; Hoffman et al., 2012). Transcription factor ChIP-seq peaks were called by each consortium for each factor using MACS using an irreproducible discovery rate (IDR) threshold of 0.05 (Zhang et al., 2008; Landt et al., 2012).

### Notation

We use the following notation below to facilitate the description of our method. We use the term “assay type” to mean a particular genomics assay protocol that may be performed in any cell type (for example “ChIP-seq targeting H3K27me3”) and “assay” to mean a particular assay type performed in a particular cell type. The term “cell type” refers to any cellular state that may be queried with a genomics assay, which may refer to any combination of cell line, tissue type, disease state (such as cancer), individual, or drug perturbation. We refer to a cell type as c and the entire set of all cell types as 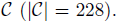. We use *a* to refer to an assay type, A for a subset of assay types, and 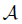 for the set of all assay types 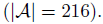. We use *s* to denote a single assay (that is, a given assay type performed in a given cell type), *S* for a set of assays, and 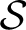 as the set of all available performed assays. Given any cell type 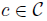 we define the set of assay types performed in this cell type as 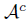 and the corresponding assays as 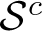. We define *I* = {1,…,*n*} as the set of all positions in a genome. An assay *s* is represented as a vector of length *n*; i.e., *s* ɛ ℝ^n^. We denote its ith entry (i.e., the value of assay *s* at genomic position *i*) as s(*i*).

### Submodular Optimization

A *submodular* function (Fujishige, 2005) is defined as follows: given a finite size *m* set *V* = {1,2,…,*m*}, a discrete set function *f*: 2^*V*^ → ℝ that offers a real value for any subset *S* ⊆ *V* is submodular if and only if:

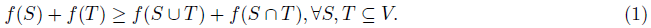

Defining 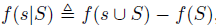), submodularity can equivalently be defined as 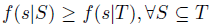 and *s* ∉ *T*. That is, the incremental gain of adding item *s* to the set decreases when the set to which *s* is added to grows from *S* to *T*. In this work, the whole set *V* represents a set of genomics assays and the set function *f*(*S*) represents a measure of quality of a subset of assays *S* ⊆ *V*.

Two other properties of set functions are relevant to this setting. First, a set function *f* is defined as *monotone non-decreasing* if

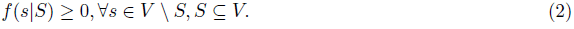

Second, we say that *f* is *normalized* if *f* (θ)=0.

In this work we are interested in the problem of maximizing a submodular function subject to a constraint on the size of the reported set. That is, we are interested in solving the problem

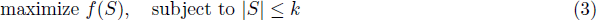

for some integer *k* ≤ |*V*|. In this work, we require that *f* is submodular, monotone-nondecreasing and normalized.

While this problem is NP-hard, it can be approximately solved by a simple greedy algorithm with a worst-case approximation factor (1 — *e*^−1^) (Nemhauser et al., 1978b). This is also the best solution obtainable in polynomial time unless *P* = *NP* (Feige, 1998). The algorithm starts with the empty set *S_0_* = θ and at each iteration *i* adds the element *s_i_* that maximizes the conditional gain *f*(*s_i_*|*S*_*i*-1_) with ties broken arbitrarily (i.e., finding *s_i_* ∊ argmax_*e*∊*V*_\*s_t_ f* (*e*|*S*_*i*−1_)) and then updates *S*_*i*_ ← *S*_*i*-1_ ∪ {*s*_*i*_}. The algorithm stops when the cardinality constraint is met with equality. This algorithm has a time complexity of *O*(*km*) function evaluations. The complexity for evaluating the facility location function is *O*(*m*^2^), if implemented naively. Since the greedy algorithm only requires computing the gain associated with adding an item to the already selected set, memorization techniques can be employed to reduce the complexity of function evaluation to only *O*(*m*), leading to an overall time complexity of *O*(*km*^2^). Furthermore, the running time can be improved to almost *O*(*m*^2^) without any performance loss by further exploiting the submodularity property (Minoux, 1978). The memory requirement of SSA depends on the choice of the optimization objective. For example, to instantiate the facility location function, one needs to compute and store a pairwise similarity graph, which takes *O*(*m*^2^) memory.

### Facility Location Function

In this work we use the *facility location* function to measure the quality of panel of assay types. The facility location function (Cornunéjols et al., 1990) *f*_fac_: 2^*V*^ → ℝ is defined as follows:

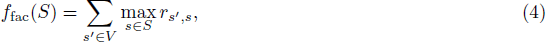

where *r*_*s*’_,_*s*_ measures the pairwise similarity between assays *s*’ and *s* (defined below). Intuitively, the facility-location function takes a high value when every assay in *V* has at least one similar representative in *S*.

### Assay Type Similarity

We use the following strategy to define the similarity between each pair of assay types in order to use this similarity to define a facility location function. We define this similarity differently depending on the application: In the selection of past assays setting, the particular assays performed in the cell type of interest c are available, while in the selection of future assays setting we must estimate this similarity from reference cell types.

In the selection of past assays setting, we directly use the signal vectors *s_i_* and *s_j_*. to derive the similarity. We define this similarity as *r_s_i_,s_j__*.=|*ρ*_*s_i_,s_j_*_ *ϵ* [0,1], where *ρ_*s_i_,s_j_*_.* is the Pearson correlation between the signal vector *S_i_* and *S_j_*. Pearson correlation is frequently used to evaluate the similarity between genomics assays (ENCODE Project Consortium, 2012). For efficiency, we compute the correlation measure *ρ_s_i_,s_j__* only across a subset of genomic positions *I’ ⊆ I*, where *I’* is randomly subsampled from *I*, and |*I*’| ≈ 0.01|*I*|.

In the selection of future assays setting, the assays in the cell type c are not available, but the assays performed in cell types other than *c*, 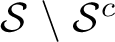, are available. Let 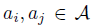 be the assay types associated with the assay *s_i_* and *S_j_*, respectively. Let 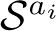 be the set of assays in 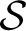 with type *a*_*i*_. We approximate the similarity between *s_i_* and *S_j_* by aggregating the all similarity between the pairs in 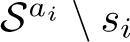 and 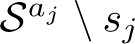. We utilize the aggregation strategy by taking the average of these similarity scores. Mathematically, the aggregated similarity is defined below:

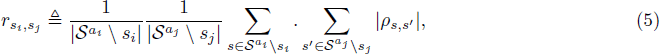

We chose to use the average correlation r because the facility location function defined via the similarities aggregated in this way correlated best with our evaluation metrics. We provide the comparison of such aggregation strategy against other strategies in the Supplementary Note 2.

### Evaluation Cross-Validation Strategy

We would prefer to apply our method once to select a single panel of assay types. However, doing so could result in a panel of assay types that have not been performed in any cell type (or very few cell types), which would prohibit evaluating the quality of this panel. Therefore, we apply a cross-validation strategy that repeatedly holds out a subset of assay types for evaluation and selects a panel from the remaining assay types, and we perform this cross-validation separately for each cell type in turn (Figure 3D). To evaluate the quality of our method with respect to a cell type *c*, we restrict ourselves to selecting from the set of assays performed in *c* (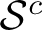). We randomly partition 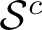 into 10 equally-sized, disjoint folds. Of the 10 folds, a single fold is retained as the target set 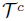 and the remaining 9 blocks are used as the source set 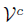. We select a panel of assays 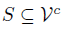 from the source set 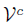 and evaluate the panel on the assays relative to the target set 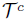 using the three evaluation metrics described below. The process is then repeated ten times, with each of the ten folds used once as the target set. We average the ten results are averaged to produce a single number representing the performance.

### Assay Imputation

The *assay imputation* evaluation metric measures the ability of a panel of assay types to be used to impute the results of other assay types outside the panel (Figure 3A). We formalize assay imputation metric as a regression problem in which the assays in the panel *S* are used as features to predict the target set assays, 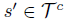. In this regression problem we have one labeled example for each position in the genome.

As our regression model, we use support vector regression with a Gaussian kernel. To construct the training and test data, we randomly choose disjoint sets of genomic positions *I^Tr^*, *I^Te^* ⊆ *I*, where *I^Tr^* ∩ *I^Te^* = θ. In our experiments, we set |*I^Tr^*|= 5,000 and |*I^Te^*|= 2,000 Given the panel *S*={*si*,…, *S*_|*S*|_}, a target assay 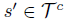, and the training genomic positions *I^Tr^*, we create the training data as 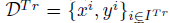, where 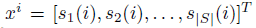 and *y^i^* = *s*’(*i*). Similarly, the test data set is constructed as 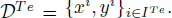. The hyperparameters of the regression model are tuned using 5-fold cross validation. We measure the performance of the trained model on the test data 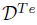 as the squared correlation coefficient θ _*s*’_. We repeat this evaluation process for every target assay in 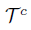 and report the performance of the panel *S* as the average squared correlation coefficient 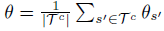.

### Functional Element Prediction

The *functional element prediction* evaluation metric evaluates how well a panel of assays can predict the genomic locations of functional elements such as promoters, enhancers and insulators. Because there are few validated examples of each type of element, we use experimentally-determined binding of transcription factors, as determined by transcription factor ChIP-seq peaks, as a proxy for functional elements. Most known types of functional elements can be characterized by the binding of particular transcription factors (Visel et al., 2009; Burgess-Beusse et al., 2002). Note that functional element prediction is similar to assay imputation in the sense that both evaluation metrics aim to predict the output of a genomics assay; however, functional element prediction focuses on just transcription factor binding sites, whereas assay imputation focuses on the whole genome. Similar to assay imputation, we consider this metric separately for each cell type. For an evaluation cell type *c*, we denote the set of transcription factor ChIP-seq assays performed in *c* as 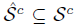. Given a bi-partition of 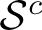 into the source set 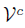 and the target set 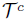, we choose from the source set 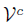 a panel of assays, and we evaluate functional element prediction only on the target assays in the set 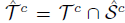, in contrast to the assay imputation metric where all assays in 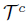 are used for evaluation.

For a target transcription factor assay 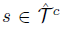, let p be a binary vector {0,1}^n^ indicating the genomic positions where *s* has a peak as called by the peak-calling algorithm. That is, *p*(*i*) = 1 if there is a peak at position *i*, and *p*(*i*)=0 otherwise. We use a support vector machine (SVM) with Gaussian kernel to predict p given a panel of assays 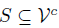. For a given testing factor p, we refer to the positions where *p*=1 as *I*_+_ and the set of positions where *p* = 0 as *I*_−_;=*I* \ *I*_+_. We randomly choose 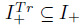 and 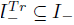 as the positive and negative positions to generate training samples. Similarly, the testing samples are randomly chosen from 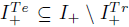 and 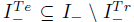. Given the panel *S* = {*s*_1_,…, *s*|_*s*_|} of assays and the set of positive training genomic positions 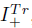, we construct the set of positive training samples as 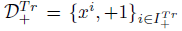 where 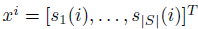. Similarly, we construct the negative training samples, positive test samples, and negative test samples as 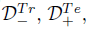, and 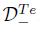, respectively. The SVM is first trained on the training data set 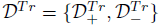, and then evaluated on the testing data set 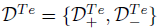.

Because there are far more genomic positions that are not a functional element than there are positions that are, measures of predictive accuracy such as the total fraction of correct predictions (“accuracy”) and the area under the receiver operating characteristic curve do not offer a reasonable measure of performance. Instead, we compute the area under the curve of a precision-recall plot (AUC-PR), which is particularly well suited for settings with imbalanced class distributions (Craven and Bockhorst, 2005; Davis and Goadrich, C). In our experiments we set 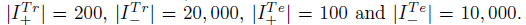 We apply 5-fold cross validation for tuning the hyperparameters of the SVM. Let *γ*_*s*’_ be the normalized area under curve for the precision-recall plot (i.e., *γ*_*s*’_ ϵ [0,1]) for each target assay 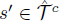 We illustrate this procedure schematically in (Figure 3B). We report the performance as the average AUC-PR on all target assays, i.e., 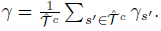.

### Annotation-Based Evaluation

The *annotation-based evaluation* metric measures the quality of a panel of genomics assays according to the quality of the genome annotation that is obtained by inputting the panel into a semi-automated genome annotation (SAGA) algorithm. SAGA algorithms are widely used to jointly model diverse genomics data sets. These algorithms take as input a panel of genomics assays and simultaneously partition the genome and label each segment with an integer such that positions with the same label have similar patterns of activity. These algorithms are considered “semi-automated” because a human performs a functional interpretation
 of the labels after the annotation process. Examples of SAGA methods include HMMSeg (Day et al., 2007), ChromHMM (Ernst and Kellis, 2010), Segway (Hoffman et al., 2012) and others (Thurman et al., 2007; Lian et al., 2008; Filion et al., 2010). These genome annotation algorithms have had great success in interpreting genomics data and have been shown to recapitulate known functional elements including genes, promoters and enhancers. We use the SAGA method Segway in this work.

In order to apply annotation-based evaluation to a panel of assays, we input this panel into a SAGA algorithm and evaluate the resulting annotation (Figure 3C). Intuitively, a diverse panel of assays input to a SAGA algorithm should more accurately capture important biological phenomena than a redundant panel.

To evaluate the quality of an annotation relative to a particular genomics data set, we use the variance explained measure (Libbrecht et al., 2015). Given an evaluation cell type *c* we randomly partition 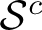 into a source set 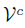 and a target set 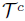 For a given panel of assays 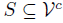 we First train a Segway model based on the panel and then obtain an annotation *y*. Segway outputs an annotation 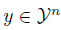 where 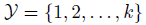 is a set of *k* labels that an annotation can take on at each genomic position. For each target assay 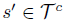 we measure the quality of the annotation *y* as how well it explains the variance of the assay *s’*. We First compute the signal mean of *s’* over the positions assigned a given label *l* as

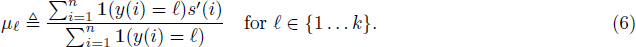

We then define a predicted signal vector *ŝ’* with *ŝ’*(*i*)= *μ*_*y*(*i*)_ and compute the prediction error as *d*_*i*_= *ŝ’*(*i*) − *s’*(*i*). We compute the residual standard deviation of the signal vector as

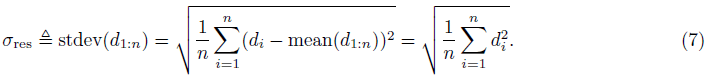

The last equality holds because mean(*d*_1:*n*_)= 0 by construction. *σ*_res_ measures the residual standard deviation of the target assay *s*’ accounting for the annotation *y*. Let *σ*_*OV*_= stdev(*s*’(1:*n*)) be the overall standard deviation of the assay *s*’ The normalized variance explained by the annotation *y* is then

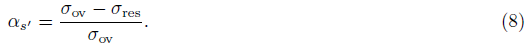

Observe that *σ*_ov_ always upper bounds *σ*_res_. The measure *α*_*s*’_ϵ[0,1] represents the fraction of the variance of the assay *s*’ explained by the annotation *y*, where larger values indicate better agreement.

In our experiments, we trained the Segway model with 10 EM random initializations (using GMTK (Bilmes and Rogers, 2015) and 15 labels at 100 base-pair resolution. We report the performance as the averaged measure on all target assays as 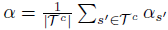.

## Source Code

SSA is supported in the form of web server available at http://melodi.ee.washington.edu/assay_panel_selection.html. Source code for SSA and the pre-computed assay type similarity matrix are also available online at http://github.com/melodi-lab/submodular-selection-of-assays.

## Author Contributions

All authors devised the method and designed the analysis. KW and MWL analyzed the data, implemented the method, developed the software and wrote the initial manuscript. All authors edited and approved the final manuscript. WSN and JAB jointly supervised the project.

## Acknowledgments

This work was supported by NIH awards R01 CA180777 and U41 HG007000.

